# Metabolic modulation in liver by glucan from *Enterococcus hirae* (OL616073) *via* gut-liver axis culture model of steatosis

**DOI:** 10.1101/2025.10.29.685250

**Authors:** Uma Rani Potunuru, Neha Gupta, Swati Tiwari, Irshad Ahmad Shah, Devendra Kumar Patel, Prathapkumar Halady Shetty

## Abstract

Metabolic dysfunction associated fatty liver disease (MAFLD) starts with increased fat deposition called steatosis progressing towards steatohepatitis, fibrosis, cirrhosis and end stage liver disease. Dysfunctional metabolism in type-2 diabetes or obesity is associated with gut dysbiosis, causing release of gut microbial endotoxins like lipopolysaccharides (LPS) into the liver via portal circulation also known as gut-liver axis. Endotoxins or LPS signals metabolic changes in the liver initiating steatosis via gut-liver axis. Various pre-clinical models have been studied to evaluate the role of fermented food derived bioactive compounds preventing the process of steatosis. Among them exopolysaccharides (EPS), as a part of fermented foods have been reported to produce metabolites by the gut microbial fermentation, providing liver with health benefits. In the present study, we have evaluated the role of a noble EPS or G (glucan) in the condition of liver steatosis by using cell culture model of gut-liver axis by co-culturing Caco-2 and HepG2 cells in transwell culture dishes. LPS was used to mimic gut dysbiosis, with or without pre-treatment of G to caco-2 cells in the upper chamber, wherease, HepG2 cells in lower chamber were extracted for metabolites to be analyzed by LC-Q-TOF. The global metabolomic profiles of LPS+G+ and LPS+ treated cells were compared. The primary target of G showed its role in cholesterol and primary bile acid metabolism along with changes in nucleotide, vitamins and branched chain aminoacid metabolites helpful in reducing fatty liver. Future studies with this glucan using targeted metabolomics could confirm biomarkers of therapeutic intervention of steatosis and could be translated into healthy food products.

## 1. Introduction

Fatty liver or steatosis is defined as fat deposition exceeding 5% of liver cells ^1^ without any inflammation. By gradual increase in fat deposition and derangement in liver metabolism, simple steatosis progresses towards hepatic inflammation or steatohepatitis, fibrosis, cirrhosis finally resulting in liver failure (Antunes C, *et. al.* 2025; ^2^. In the natural history of fatty liver diseases, steatosis or non-alcoholic fatty liver disease (NAFLD), recently termed as metabolic dysfunction associated fatty liver disease (MAFLD) ^3, 4^ advances to end stage non-alcoholic steatohepatitis (NASH) or metabolism associated steatohepatitis (MASH). The pathogenesis of MAFLD is mainly associated with metabolic dysregulation causing changes in gut microbiome also known as gut-dysbiosis, releasing micorbial metabolites and toxins to be transported to the liver via gut-liver axis, that signals the changes in hepatic cells ^5^. There are evidence of gut microbiome signaling the organ function outside gut termed as gut-organ axis ^6–8^. One such axis is gut-liver axis which is formed by the close anatomical and functional bidirectional interaction through portal circulation. In metabolism associated gut dysbiosis, release of microbial endotoxin lipopolysaccharide (LPS) into the liver by portal vein, signals the dysregulated lipid metabolism or lipid accumulation (lipotoxicity), anti-oxidant imbalance (oxidative stress) and inflammation in the liver cells (^9, 10^. The global incidences of fatty liver diseases are on the rise due to lifestyle and environmental risk factors ^11^. Currently, no pharmacological treatments have been approved to prevent its outbreak. Thus, basic and preclinical research becomes most essential to improve our understanding of pathophysiology and the development of interventional therapeutics. Among the various *in vitro* cell culture models of NAFLD, human hepatocyte cell line monolayer 2D culture or co-culture with liver parenchyma have been reported ^12^. Though the immortalized monolayer cell culture model is stable ^13, 14^ and could provide reproducible results with simplicity of handling, these models limit the complete pathophysiology of gut dysbiosis-mediated liver diseases. Thus, in the present study, we used a co-culture of monolayer cell lines of Caco-2 and HepG2 in a transwell culture dish to mimic the gut-liver axis and used LPS as an agonist to mimic gut dysbiosis^15^.

Fermented foods and their effects on gut health have been reported, and the underlying mechanism of this effect is attributed to probiotics (a class of beneficial microbes) and their derived metabolites in the form of bioactive compounds ^16^. One such class of bioactive compounds are exopolysaccharides (EPS), polymers of sugars produced by microorganisms, which are also part of the human gut microbiome, via the process of natural biological fermentation of the carbohydrates. These EPSs have recently termed as released postbiotics. These released postbiotics, when exposed to gut microbe metabolic pathways, produce soluble factors termed as metabolic postbiotics such as short chain fatty acids (SCFAs), polyamines, tryptophan derivatives, hydrogen sulphide, polyphenol metabolites, etc., based on their origin ^17^. Several studies have reported these metabolic postbiotics to be promising in the therapeutic intervention of MASLD ^18^.

In the present study, we aim to understand the role of *in vitro* fecal fermented EPS (characterized as glucan-a homopolysaccharide of glucose units) on liver cell metabolism in steatosis conditions, in a co-culture model of gut-liver axis. *In vitro* fecal fermented EPS was used for testing, as fecal microbial fermentation of EPS is known to release metabolites (metabolic postbiotics/ soluble factors) of therapeutic interest ^19, 20^. To achieve this objective, we standardized an *in vitro* liver steatosis model by co-culturing Caco-2 (colon cancer cell line to mimic gut epithelium) and HepG2 (liver cell line) in a transwell culture dish. Lipopolysaccharide (LPS), an endotoxin produced by predominant gram-negative bacteria in dysbiosis state of the gut ^15^, was used as a stimulant on caco-2 in the upper chamber to mimic gut dysbiosis; with HepG2 co-cultured in the lower chamber of the transwell culture dish, modeling gut-liver axis. Lipid accumulation and fat metabolic pathway gene expression changes were evaluated by oil red O staining and quantitative RT-PCR, respectively, validating the condition of steatosis. Further, glucan was pretreated with LPS post-challenged co-culture to evaluate the differential metabolic profiles by employing high-resolution mass spectrometry (LC-MS/MS) on the extracted metabolites of HepG2 cells. The global metabolic data analysis of liver cell metabolites revealed differential metabolome profiles between glucan-treated and untreated LPS-challenged liver cells with regard to lipid, amino acid and vitamin metabolic pathways, as well as primary bile acid biosynthesis pathways in liver cell. A thorough literature analysis of these metabolites was carried out to understand the role of glucan on changes in molecular dynamics involved in steatosis, and the potential role in protecting the liver from lipotoxicity, inflammation, and cellular oxidative stress. Thus, the current study shows the metabolome shift upon glucan exposure to fatty liver cells, thereby providing insight into its therapeutic potential in preventing hepatic injury.

## 2. Materials and Methods

### 2.1 Materials

Human intestinal cell line-(Caco-2, NCCS, Pune, India), Human liver cancer cell line-(HepG2, NCCS, Pune, India), Dulbecco’s modified Eagle’s medium (DMEM; Cat#AL007S), Minimum Essential Media (MEM; Cat#AL047S), Fetal Bovine Serum (FBS; Cat#RM9955), Antibiotic antimycotic Solution 100X Liquid (w/10, 000 U Penicillin, 10mg Streptomycin and 25 μg Amphotericin B per ml in 0.9% normal saline, Trypsin-EDTA solution) (Cat#A002), MTT (3-(4, 5-dimethylthazolk-2-yl)-2, 5-diphenyl tetrazolium bromide) (Cat#TC191), Phosphate buffered saline (PBS, Cat# TL1101) were from Himedia, India; Dimethyl sulphoxide (DMSO) (Cat# SE9S690359) was from Merck, India. Corning® Transwell® polyester membrane cell culture inserts 12 mm Transwell with 0.4 μm pore polyester membrane insert, TC-treated, sterile, 48/cs - 48EA (CLS3460-48EA); Lipopolysaccharide from Escherichia coli O111:B4 (L3023). Other materials used for this study are mentioned in respective methods section.

### 2.2. Fermented exopolysaccharide (EPS/Glucan) preparation

Fermented EPS**/**glucan (or G) was prepared as per the procedure standardized in our lab. Briefly, Glucan producing lactic acid bacteria, which were isolated from an Indian traditional fermented food (Fourteen hours fermented idli batter) and stored as glycerol stock in our lab, were streaked on De Man, Rogosa and Sharpe (MRS) agar plates with and without 2 % sucrose. Colony morphology, Gram character, and catalase test culture were studied. Further, this EPS producing culture was maintained in MRS broth complemented with 40 % glycerol at −80 ℃. Species-level identification was done by 16S rRNA gene analysis method. EPS isolation was carried out according to the procedure described by Kavitake *et al*. ( ^21^. After the dialysis against Milli-Q water using a 12–14 kDa Molecular weight cut-off (MWCO) dialysis membrane at 4 °C for two days, the dialyzed EPS was lyophilized. Further, the lyophilized powder form was purified by using an anion exchange DEAE Cellulose-52 column and a size exclusion Sephacryl S-300 column. Characterization was done as per the standardized methods carried out by the PI’s group mentioned by Kavitake *et. al*.^21^. It is characterized as a glucan EPS.

For fecal fermentation, fresh human fecal samples were obtained from healthy human volunteers (two females and two males, subjects aged between 25 and 52 years with no history of antibiotics intake for the past three months, and were free from any kind of intestinal disorders). This study was approved by ‘Institutional Ethical Committee (Human Studies) with the approval number HECPU/2021/30/07-12-2021. Samples were prepared as per the procedure standardized in our lab. Samples were homogenized in saline and made into 10% (v/v) fecal slurry and centrifuged to remove large particle matter at 500g for 5min and used as fecal inoculum. The growth medium was prepared according to the protocol described by Mengke yao *et.al.*^22^ and slightly modified according to Wang *et. al*. and Jia *et.al.* ^19, 23^. The medium was prepared by mixing yeast extract (2.0 g), peptone (2.0 g), bile salts (0.5 g), L-cysteine (0.5 g), NaHCO_3_ (2.0 g), NaCl (0.1 g), hemin (0.02 g), CaCl_2_⋅6H_2_O (0.01 g), MgSO_4_⋅7H_2_O (0.01 g), K_2_HPO_4_ (0.04 g), KH_2_PO_4_ (0.04 g), vitamin K_1_ (0.01 g), resazurin (0.01 g), and Tween-80 (2.0 mL), in 1.0 L distilled water. After the pH was adjusted to 7.6 ± 0.1, the fermentation medium was autoclaved at 121 °C for 20 min and divided into sterile anaerobic vessels. EPS was added at the rate of 1% in fermentation medium (FM) mixed with fecal inoculum (FI) (FM: FI=9:1) and incubated at 37 °C for fermentation. The three replicates of the 24-hour fermented EPS were pooled together to make 100 mg equivalent of unfermented EPS in sterile cell culture media and filtered through 0.45µm PES sterile filters (Millex 33mm-SLHP033RS Merck) for use in cell culture experiments. The purification and functional characterization of the glucan was previously published by our group ^21^. Further prebiotic characterization of the fermented glucan was carried out by our group and had confirmed short chain fatty acid (SCFAs) production with potential health benefits (Swati Tiwari *et. al.*‘Evaluating the prebiotic potential of a glucan EPS from Enterococcus hirae OL616073: Digestive resistance, probiotic growth stimulation, and gut microbiome modulation’, Food Research International, vol.222, part 1, December 2025, 117649).

### 2.3 Cytotoxicity analysis

The effect of fermented glucan on cell viability was determined by MTT assay as per standardized protocol as described by Kumar *et. al.* ^24^. Briefly, Caco-2 and HepG2 cells were seeded at a density of 10, 000 cells/well in sterile 96-well cell culture plates in MEM medium with 10 % FBS and incubated at 37°C for 24 hours under a humidified atmosphere with 5 % CO2. After 24 hours, cells were exposed to serum-free media overnight for synchronization. Filter sterilized fermented EPS at concentrations of 1.25, 2.5, 5, 10, and 20mg/ml (mg weight equivalent of unfermented EPS) were treated to Caco-2 cells for 24 hours, followed by removal of cell-free supernatant and treating the supernatant to HepG2 cells cultured simultaneously in another 96-well plate. At the end of 24 hours’ treatment, culture media from HepG2 cells were removed, and rinsed with PBS. 200 μL of MTT solution (0.5 mg/mL) was added to each well except the blank and incubated at 37 °C for 3.5 hours to 4 hours, following which MTT solution was removed and 200 μL of DMSO was added, and the color developed was measured at 570 nm using an Eion plate reader (Biotek, India).

### 2.4 Cell culture and co-culture design

Both Caco-2 and HepG2 cells were maintained in MEM with 10 % fetal bovine serum (FBS) and 1X antibiotic-antimycotic solution, at 37 °C with 5 % CO2. The Caco-2 and HepG2 cells were sub-passaged with Trypsin-EDTA. All experiments with Caco-2 were carried out between passage no. 40 to 60, whereas HepG2 cells were between passage no. 3 to 15. The monolayer of Caco-2 cells and HepG2 was visualized under the inverted phase contrast microscope (Nikon). A monolayer of Caco-2 cells was cultured in serum-free culture media on the trans-well filter inserts for 20±1 days. The lower chamber of the 12-well plate with inserts was layered with HepG2 cells, a week prior to the treatment, obtaining a monolayer, mimicking the gut-liver axis.

### 2.5 Steatosis model

Caco-2 cells in the upper chamber of the transwell plate were treated with or without LPS at 10µg/mL of concentration for 24 hours, with HepG2 cells in the lower chamber of transwell inserts, seeded at 60 to 70% confluence. At the end of 24 hourstreatment period media was removed from both chambers and HepG2 cells were analyzed for increased fat deposition and altered fat metabolism by oil red O staining and gene expression by quantitative RT-PCR. Cells without any treatment act as a negative control.

#### 2.5.1 Oil Red O staining for fat accumulation

Oil Red O staining was carried out as per a modified method referred from Hyeon-Soo Jeong *et. al.* and Meng-ya Shan *et. al.* ^25, 26^. Briefly, at the end of treatment, media was removed and cells were rinsed with ice-cold 1XPBS and fixed with 10%formalin. Fixed cells were incubated with the diluted Oil Red O (0.3%) at room temperature for 2 hr. At the end of the incubation period, cells were washed with sterile distilled water, and fat droplets in the stained cells were visualized under light microscopy and photographed using a Nikon inverted microscope TS-100F. For quantitative analysis of Oil Red O content levels, isopropanol was added to each sample and then shaken at room temperature for 5 min. The absorbance of the samples was read at 510 nm using an Eion microplate reader (Biotek, India).

#### 2.5.2 Fat metabolism gene expression by RT-PCR

HepG2 cells from the co-culture experiment were subjected to RNA isolation as per the manual method using TRI reagent (Cat#MB601, Himedia, India) and chloroform (Cat#MB106, Himedia, India), and concentration was determined by NANO-DROP LITE (Thermo-scientific)immediately converted to cDNA using Verso cDNA kit (Cat#AB1453, Thermo-scientific, India) as per the manufacturer’s instructions, RT-PCR was set up in Eppendorf. Further, Real Time PCR was carried out using Takara’s TB Green® Premix Ex Taq ™ (Tli RNaseH Plus) kit as per the manufacturer’s protocol, and PCR was set up in Insta Q48 M2 (Hi-Media) instrument. The key human lipid metabolism pathway genes reported to be altered in fatty liver diseases ^15, 25, 27, 28^ were targeted viz. Acetyl CoA carboxylase beta (ACCB), 3-hydroxy-3-methylglutaryl-CoA reductase (HMGCR), Fatty acid synthase (FAS), Sterol regulatory element binding protein (SREBP), Patatin-like phospholipase domain containing 3 (PNPLA3), and Peroxisome proliferator-activated receptor alpha (PPARα). Oligos for primers were ordered from Eurofins Scientific, India. The mRNA expression was normalized with reference gene human β-actin (housekeeping gene). The relative quantification of mRNA level was determined using the 2−ΔΔCt method.

All primers are listed in Supplementary Table 1.

### 2.6 Metabolite profiling

#### 2.6.1 Cell culture treatment

Cells were treated with or without LPS (10µg/ml) as mentioned in section 2.5 for steatosis model, in addition to glucan pre-treatment at 10mg/ml (as there was 100% cell viability up to 20 mg/ml, half of the maximum concentration i.e.10 mg/ml was used for metabolome analysis) for 24 hours in LPS challenged condition. Three independent experiments were carried out to obtain three biological replicate samples.

#### 2.6.2 Intracellular metabolite extraction and sample preparation-

Metabolite extraction and sample for the LC-Q-TOF study were prepared as per a modified method described by Zhang *et al.* ^29^. Briefly, at the end of the treatment, the media was removed from both upper and lower chambers of transwell dish and HepG2 cells in lower chamber, were rinsed with ice-cold 1XPBS twice. An ice-cold solution of methanol and water at a 80:20 ratio was added to the wells, and HepG2 cells were scraped and transferred to a fresh autoclaved 1.5 ml micro-centrifuge tube. The tubes with cells in the methanol and water mix were subjected to three cycles of freeze (in liquid nitrogen) and thawing (at room temperature) in order to disrupt the cell membrane, followed by centrifugation at 12000 rpm for 20 min at 4 °C. The supernatant samples now contain the extracted intracellular metabolites and were transferred to sterile tubes and air dried in a sterile cabinet to be stored at −80°C till further processing.

#### 2.6.3 Analysis by LC-Q-TOF-

##### LC Conditions

Non-targeted metabolomics analysis was performed on a total of 9 samples by LC-Q-TOF. Liquid chromatographic analysis was done on the UPLC machine AB SCIEX Exion LC 2.0 system (AB SCIEX, Foster City, CA, USA). For the analysis of metabolites, a Kinetex 1.7 µm EVO C18 100 Å (100 x 2.1 mm, 1.8 µm) LC column was employed. Kinetex 1.7 µm EVO C18 100 Å was found to have the best performance for the untargeted metabolites screening. The binary solvent system comprising of 10 mmol ammonium acetate in 0.1% acetic acid in milli Q (Solvent A) and 10 mmol ammonium acetate in 0.1% acetic acid in 5% water with 95% acetonitrile (Solvent B). The total run time of the gradient flow was 20 minutes, with a flow rate of 0.400 mL/min. Initially, B was kept at 5 %, increasing to 100 % at 15 minutes, and kept for 18.5 minutes. B was then decreased to 5 % at 19.5 minutes and kept at 5 % for 20 minutes. The volume of each sample injected into the LC system was adjusted to 5.0 µL. The illumination on the sample manager was turned off to prevent any additional degradation of the sample, and the temperature of the sample manager was kept at 4 °C to prevent any loss of analyte. The temperature of the column was kept at 40 °C to allow clear separation of the metabolites on the column.

##### Mass conditions

The AB SCIEX LC-Q-TOF 5600+ system (AB SCIEX, Foster City, CA, USA) was operated using an electrospray ionization (ESI) probe with the Turbo V ion source. The source parameter for mass analysis are GS1 (40.0 psi), GS2 (40.0 psi), CUR (30.0 psi), TEM (400 ℃), ISVF (5500 V), TOF Mass range (50 to 1500 Da), and Accumulation Time (0.025 sec). The TOF MS scan parameters in the positive mode were declustering potential (100 eV) and collision energy (10 eV). The data was obtained utilizing the IDA (Information dependent acquisition) scan method. The MS mass range studied was 50-1500 m/z using optimum source conditions, and the MS/MS was obtained using a mass range of 50-1250 m/z and a 0.025 sec accumulation period.

#### 2.6.3 Data processing

The raw files obtained after AB SCIEX LC-Q-TOF analysis were converted into mzXML files using ProteoWizard. The mass spectral data and peak picking were performed using Metaboanalyst (v 5.0). The program detects and identifies endogenous metabolites that differ between samples by integrating raw data processing, retention time correction, and statistical analysis.

### 2.7 Statistical analysis

Student’s t-test and One Way ANOVA were performed to compare the means of two or more groups in cell culture and Real time PCR experiments. For metabolomics analysis, univariate and multivariate data analysis were performed by the Statistical Analysis (one factor) module of Metabo-Analyst 5.0. For other purposes, SPSS 16.0 was applied for one-way ANOVA (analysis of variance) using Student t-test.

## 3. Results and discussion

### 3.1 Cytotoxicity of fermented glucan

To determine the cytotoxicity of fermented glucan towards HepG2, differentiated Caco-2 cells were incubated with 0, 1.25, 2.5, 5, 10 and 20 mg/ml (weight equivalent of unfermented glucan) concentrations of fermented glucan in for 24 hours, followed by the assessment of cell viability of HepG2 cells as described in the method section 2.3. The data showed no cytotoxicity expressed in percentage cell viability (100%) of HepG2 cells following treatment up to 20 mg/ml concentrations of glucan (result is shown in Supplementary Figure S1).

### 3.2 Steatosis model in co-culture of Caco-2/HepG2 mimicking gut-liver axis

We evaluated the deposition of fat as a functional establishment of model of fatty liver or steatosis by treating Caco-2 with pathological doses of lipopolysaccharide (10µg/ml) which can cause pathological breach in caco-2, and can leak in to HepG2 cells via transmembrane filter to elicit liver pathogenesis in the form of steatosis or steatohepatitis. There was significantly increased deposition of oil droplets in LPS-treated HepG2 cells compared to no treatment control (Figure 1a). Thus, we further determined changes in liver fat metabolism upon LPS treatment to Caco-2/HepG2 co-culture through changes in genes involved in the fat metabolic pathway in HepG2 by quantitative RT-PCR analysis. Among the screened fat metabolism genes, such as fatty acid synthase (FAS), Acetyl CoA carboxylase beta (ACCB), 3-hydroxy-3-methylglutaryl-CoA reductase (HMGCR), Sterol regulatory element binding protein (SERBP), Patatin-like phospholipase domain containing 3 (PNPLA3), and Peroxisome Proliferator-Activated Receptor alpha (PPAR-α), only ACCB and HMGCR showed significant changes upon LPS treatment as compared to untreated control. LPS treatment resulted in almost 2-fold increase and 0.5-fold reduction in HMGCR and ACCB expression, respectively, in LPS-treated HepG2 compared to no treatment control. The results of the significantly changed genes alone were shown in Figure 1B. Thus, the co-culture model of gut dysbiosis and consequent altered lipid metabolism leading to fat deposition indicated the model of steatosis in liver cells. Using this gut-liver axis steatosis model, we further tested the effect of fecal fermented glucan on liver metabolism.

**Figure 1.**
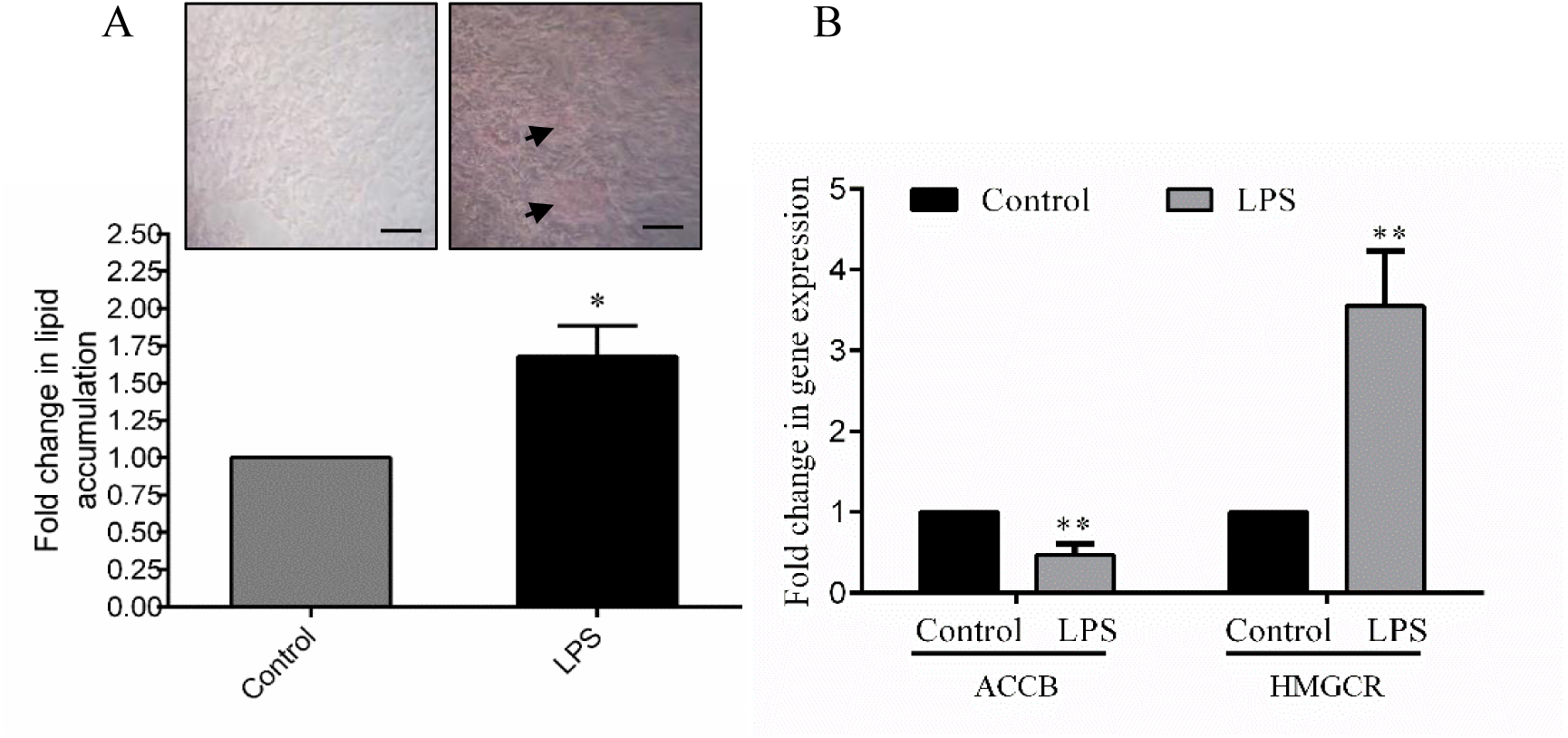
Liver steatosis markers in HepG2 cells (co-cultured with caco-2 cells). (A) Quantification of oil Red O in bargraph with representative microscopic images (10X magnification, Nikon); (B) Bar graph showing gene expression of ACCB (Acetyl Co-A carboxylase beta) and HMGCR (3-hydroxy-3-methylglutaryl-CoA reductase) in response to 24 hours LPS (10µg/ml) treatment compared to no treatment control in HepG2 cells. Error bars Mean ± SEM represent minimum of three independent experiments. \**p*<0.05, \*\**p*<0.01.

### 3.3 Comparative hepatocyte metabolome profiles in LPS and glucan challenge

A comparative metabolome profile of LPS treated HepG2 cells with or without fermented EPS (Glucan/G+) pretreatment was carried out and compared with no treatment control to understand the latter’s effect on the intrinsic dynamics of liver cell metabolism. At the end of the treatment, intracellular metabolites of HepG2 cells were extracted as per the described method in section 2.6.2 and subjected to data acquisition using LC-Q-TOF spectra in positive ion mode.

Univariate analysis showed a total of 28 significant metabolite features in the *t-test* at p<0.05 (Figure 2A and B) in both Control vs LPS+ and LPS+ vs LPS+G+ comparison profiles. Among these, only 22 metabolite features had a VIP score ≥ 1.0. Hence, fold change (FC≥1.5) analysis (figure 2C and D) of the 22 metabolites showed 11 upregulated and 11 downregulated metabolites in comparison groups of control or C vs LPS+ and LPS+ vs LPS+G+, which are listed in table 1. Changes in metabolite features observed between comparison groups is shown by the feature exclusiveness in Venn diagram (Figure 3). As the analysis was global and comparative to find exclusive and common metabolites between LPS^+^ G^-^ and LPS^+^ G^+^, we analyzed the metabolites of LPS+ with control cells (C vs LPS+) which reflected 55 LPS exclusive, 31 glucan-EPS (G+) exclusive and 234 common metabolite features indicating large number of metabolites being affected by glucan pretreatment in LPS treated HepG2 cells. Statistics and chemometric analysis were employed using defined parameters [m/z, retention time (rt), peak intensity and p value] which elucidated 14 signature metabolites to be different between control and LPS^+^ HepG2 cells (C Vs LPS+) whereas 8 metabolites were significantly different between G- and G+ of LPS^+^ HepG2 cells (LPS+ vs LPS+G+) as summarized in figure 3.

**Figure 2.**
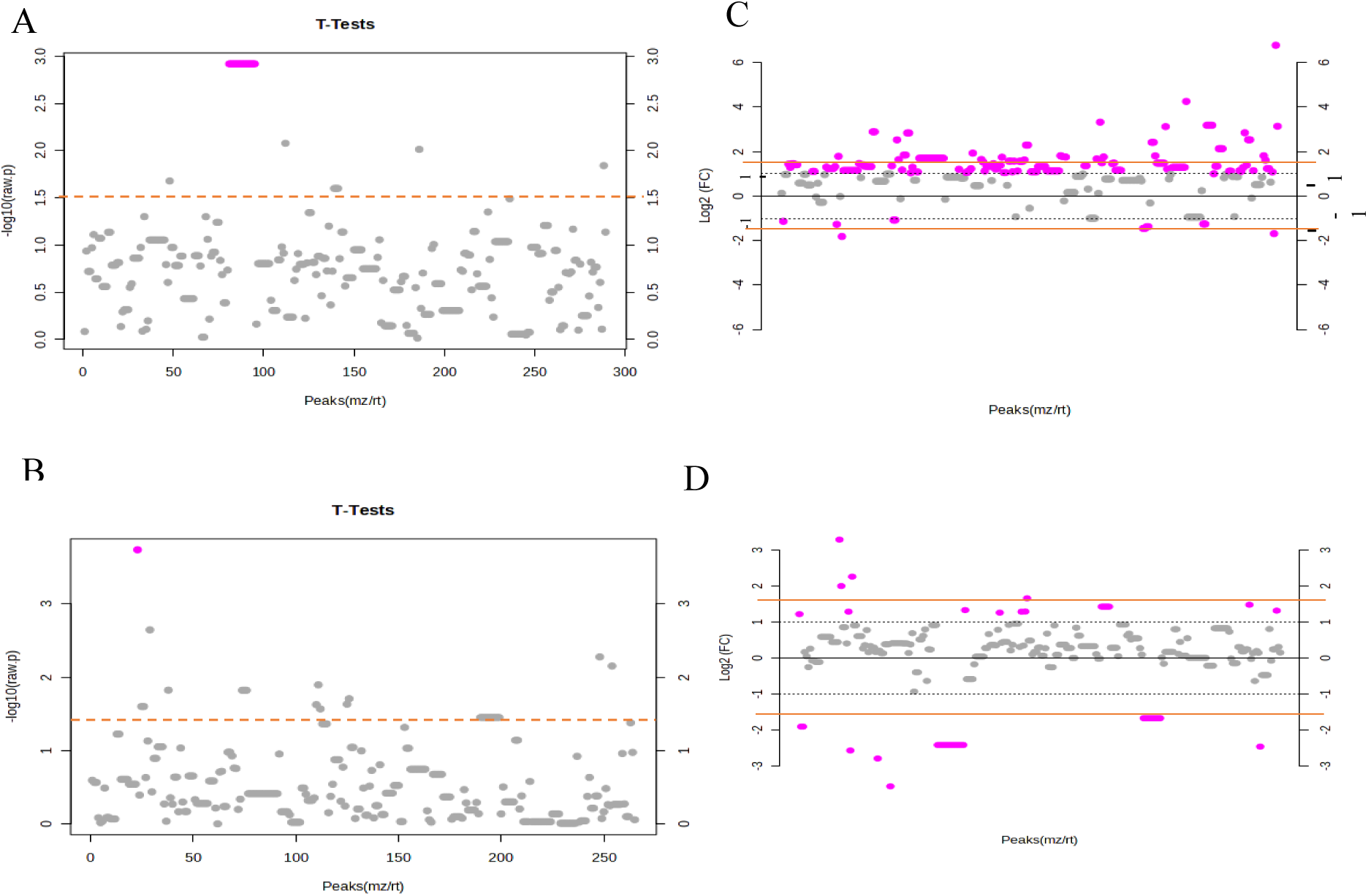
Univariate feature analysis in MataboAnalyst 5.0. Significance test in *t* test (Wilcoxon rank-sum tests) identified (A) 20 and (B) 19 significant metabolite features in Control vs LPS+ and LPS+ vs LPS+G+ comparison respectively at (*p* ≤ 0.05); Further fold change (FC threshold ≥ 1.5) analysis of metabolite features identified (C) 20 and only (D) 8 significantly up and down regulated metabolite features in Control vs LPS+ and LPS+ vs LPS+G+ cell metabolites respectively.

**Figure 3.**
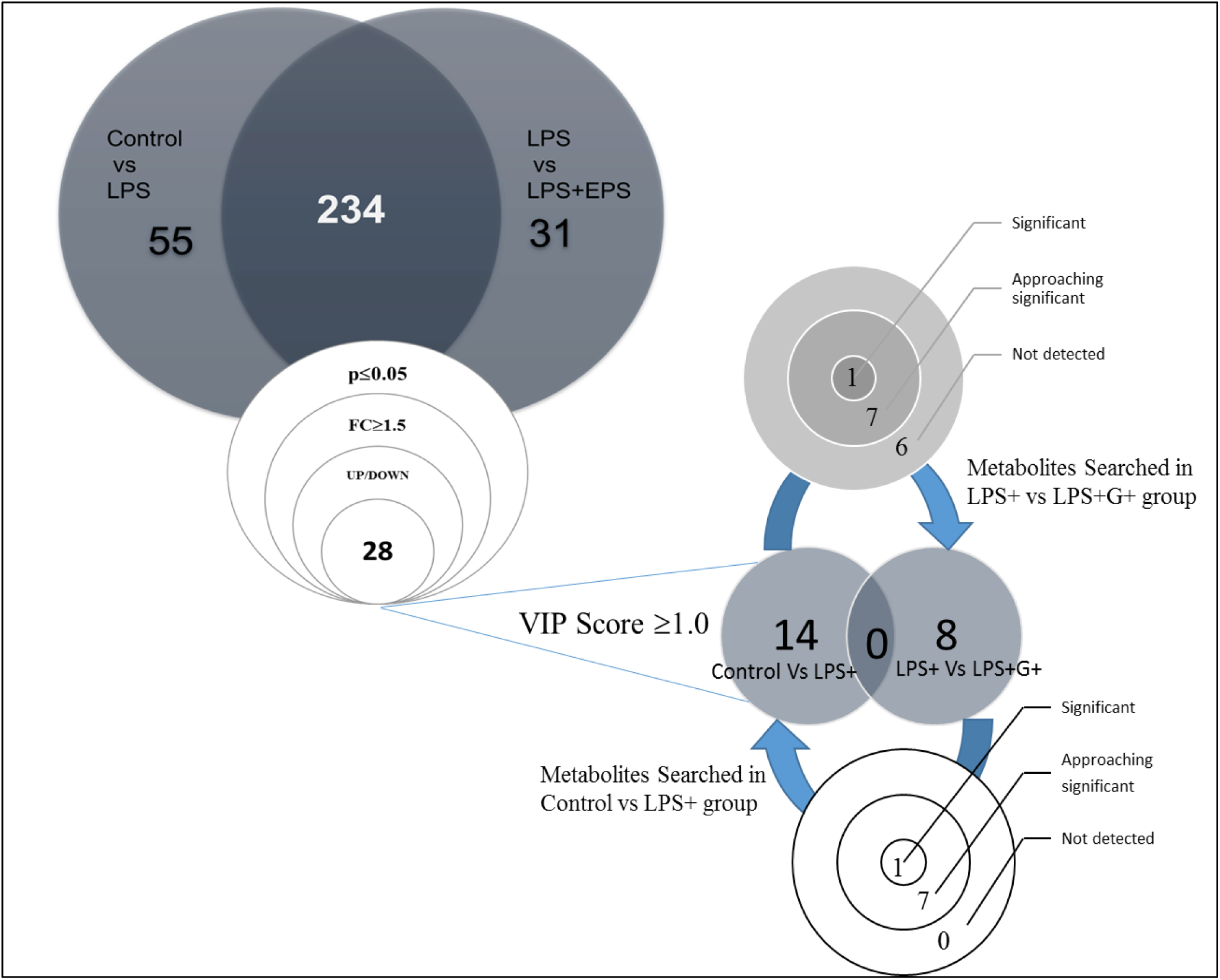
Summary of metabolite features based on statistical analysis and set threshold. Out of a total of 234 common metabolite features between two comparison groups of Control vs LPS+ & LPS+ vs LPS+G+, subjected to cutoff value of p≤0.05 and fold change ≥1.5, only 28 metabolites were obtained. Further, based on VIP score ≥1.0 the two groups had 14 and 8 exclusive metabolite features respectively.

**Table 1.** Up and down regulated putatively annotated metabolites based on significant FC (≥1.5) (p≤0.05) and VIP score (≥1.0) in glucan challenged LPS induced steatosis model. r.t. retention time, FC fold change, annotated using Human Metabolome Database (HMDB) and KEGG compound search.

| Sl No | m/z | r.t. (min) | FC value | VIP score (OPLS-DA model) | KEGG identifier | Annotation | Metabolic class |
| --- | --- | --- | --- | --- | --- | --- | --- |
| **Significantly Upregulated** |  |  |  |  |  |  |  |
| **Control vs LPS** |  |  |  |  |  |  |  |
| 1 | 346.046 | 0.655 | 2.19 | 1.23 | C00051 | Glutathione | Peptide |
| 2 | 267.059 | 0.657 | 1.66 | 1.16 | C00299 | Uridine | Nucleoside |
| 3 | 123.055 | 1.038 | 1.57 | 1.09 | C05828 | Methylimidazoleacetic acid | Imidazolyl carboxylic acid |
| 4 | 137.046 | 0.895 | 2.31 | 1.43 | C00262 | Hypoxanthine | Nucleotide |
| 5 | 203.223 | 0.802 | 2.37 | 1.66 | C00750 | Spermine | Polyamine |
| 6 | 124.039 | 0.659 | 4.21 | 1.61 | C00253 | Nicotinic acid | Pyridine |
| 7 | 124.039 | 0.659 | 4.21 | 1.67 | C03824 | 2-Aminomuconic acid semialdehyde | Alpha amino acid |
| 8 | 124.039 | 0.659 | 4.21 | 1.61 | C10164 | Picolinic acid | Pyridine |
| **LPS vs LPS+G+** |  |  |  |  |  |  |  |
| 1 | 513.989 | 6.793 | 29.05 | 1.87 | C00131 | Deoxyadenosine triphosphate | Nucleotide |
| 2 | 141.066 | 0.662 | 7.02 | 1.89 | C00153 | Niacinamide | Pyridine |
| 3 | 185.081 | 13.727 | 1.84 | 1.43 | C00588 | Phosphorylcholine | Phospholipid |
| **Significantly Downregulated** |  |  |  |  |  |  |  |
| **Control vs LPS** |  |  |  |  |  |  |  |
| 1 | 348.289 | 12.581 | 3.97 | 1.34 | C11695 | Anandamide | Lipids |
| 2 | 437.374 | 12.271 | 1.73 | 1.05 | C01753 | Beta-Sitosterol | Phytosterol |
| 3 | 437.374 | 12.271 | 1.73 | 1.05 | C04530 | 4,4-dimethyl-5alpha-cholest-7-en-3beta-ol | Steroid |
| 4 | 437.374 | 12.271 | 1.73 | 1.05 | C15915 | 4,4-Dimethyl-5a-cholesta-8-en-3b-ol | Steroid |
| 5 | 375.287 | 19.734 | 1.62 | 1.02 | C02528 | Chenodeoxycholic acid | Bile acid |
| 6 | 375.287 | 19.734 | 1.62 | 1.02 | C04483 | Deoxycholic acid | Bile acid |
| **LPS vs LPS+G+** |  |  |  |  |  |  |  |
| 1 | 132.077 | 0.652 | 7.97 | 1.74 | C00300 | Creatine | Amino acid |
| 2 | 457.113 | 3.851 | 2.12 | 1.76 | C00061 | Flavin Mononucleotide | Nucleotide |
| 3 | 198.038 | 0.665 | 1.54 | 1.50 | C02059 | Vitamin K1 | Phylloquinone |
| 4 | 198.038 | 0.665 | 1.54 | 1.50 | C04281 | Pyrroline hydroxycarboxylic acid | Amino acid |
| 5 | 198.038 | 0.665 | 1.54 | 1.50 | C04282 | 1-Pyrroline-4-hydroxy-2-carboxylate | Pyrroline |

However, there were no common metabolites between the comparison of cellular metabolites in C vs LPS+ and LPS+ vs LPS+G+, thus we explored the status of changed metabolites in C vs LPS+cells in LPS+ vs LPS+G+ metabolic features, to determine their status shift based on their statistical significance and fold change threshold.

### 3.4 Differential metabolites in LPS and glucan challenge

To understand the effect of G+ treatment on metabolite profile of HepG2 subjected to LPS mediated liver cell steatosis via gut liver axis culture model; A clear differentiation of metabolite features was shown in volcano plot for LPS+ vs LPS+G+ (figure 4A). The trends of distribution analyzed by multivariate Principle Component Analysis (PCA) model was able to highlight descriptively the trends of relationship between and within LPS+ and LPS+G+ samples for distribution in two principal components, PC1 and PC2 and LPS+ vs LPS+G+ clustering. The differentiation of metabolite features was revealed by variance of 50.4% along PC1 and 22.1% along PC2 in the PCA score plot (Figure 4B). A supervised Partial Least Squares-Discriminant Analysis (PLS-DA) model ^30^ gave a comprehensive and conclusive interpretation of the metabolic responses in LPS treated and G+ co-treated cell metabolites to govern discrimination between these two treatment groups leading to unique metabolite or biomarker identification. In PLS-DA metabolite features showed discrimination between LPS+ and LPS+G+ having covariance of 45.2% at component 1 (x-axis) and 19.3% at component 2 (y-axis) (Figure 4C). This PLS-DA model was cross-validated by analyzing predictability and non-overlapping of the model by reflecting positive Q2 (Figure 4D).

**Figure 4.**
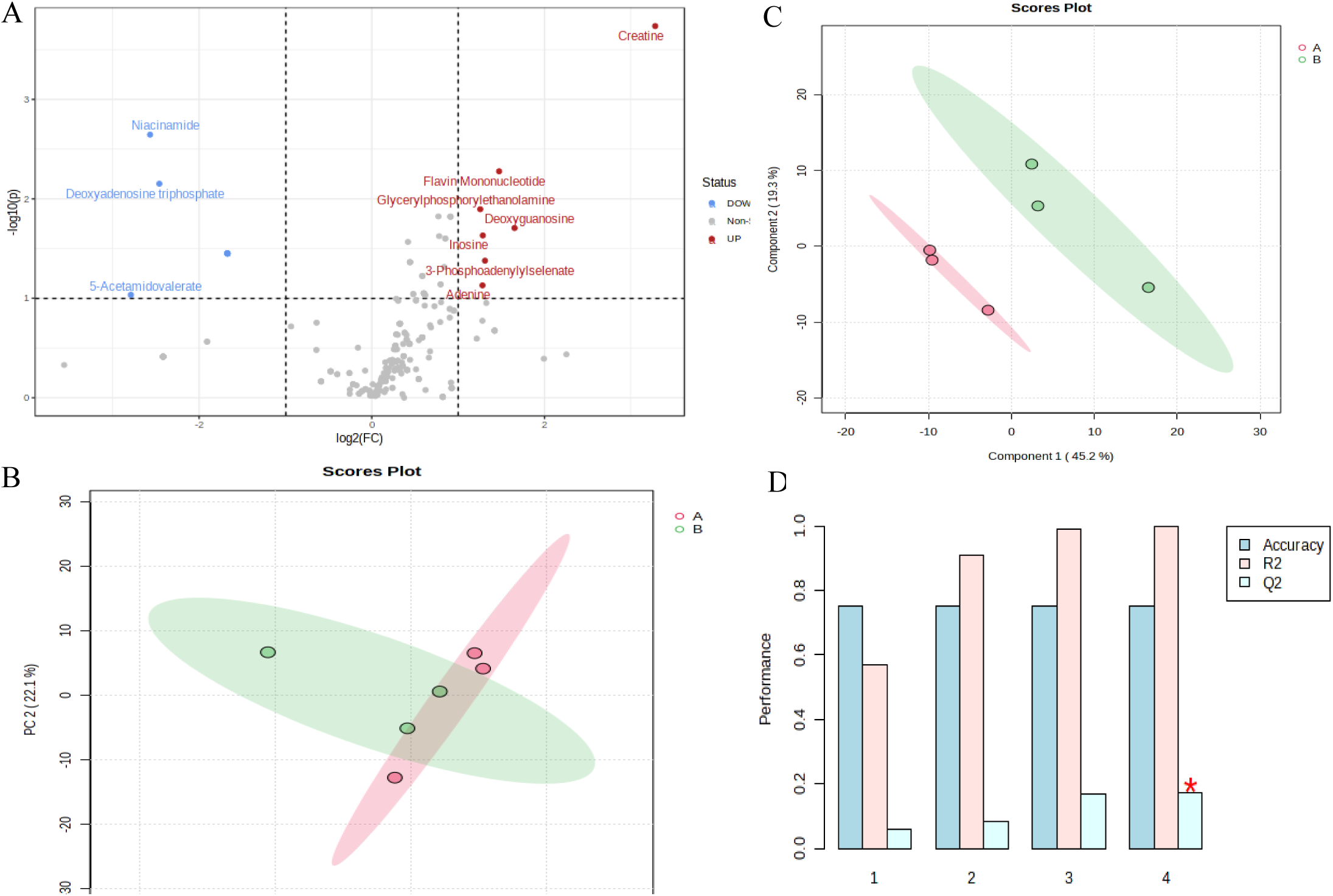
Differential features in LPS+ liver cells with or without glucan. (A) Volcano plot, showing Up- and down regulated significant (p≤ 0.05) metabolite features with clear differentiation; (B) A distinct relationship between samples and differences in LPS+ versus LPS+G+ samples along x-axis (PC1) and within groups along y-axis (PC2) are shown by principal component analysis (PCA) score plot, with a variance of 50.4% and 22.1% at 95% confidence interval by PC1 and PC2 respectively; (C) Covariance between component 1 (45.2%) at x-axis and component 2 (19.3%) at y-axis was represented by Partial Least Squares-Discriminant analysis (PLS-DA) scores plot of metabolite features in PLS-DA model; (D) A cross validation showed positive Q2 reflecting predictability and non-overlifting of the PLD-DA model.

The 14 metabolite features showing significant difference (p≤0.05) and a fold change ≥ 1.5 in C vs LPS+ were searched in cell metabolites in group of LPS+ vs LPS+G+ cells, those showing significant or approaching significance with reversed trend were considered to be unique metabolites affected in hepatic steatosis as a result of G+ treatment (Figure 3 and Table 2). These unique metabolites were further analyzed and correlated with fatty liver diseases.

**Table 2.** Differential metabolites in steatosis compared in glucan treated HepG2 cells. FC-fold change. Glutathione* showed significant change in both the comparison groups of metabolites.

| SL NO. | KEGG identifier | Annotation | Status in other comparison group |  |  |
| --- | --- | --- | --- | --- | --- |
| **Control vs LPS+** |  |  | **LPS+ vs LPS+G+** |  |  |
| Upregulated<br>$p \leq 0.05$; $FC \geq 1.5$ | | | p-value | FC | Up/Down |
| 1 | C00051 | **Glutathione** * | 0.08 | 1.44 | Down |
| 2 | C00299 | Uridine | Not detected |  |  |
| 3 | C05828 | Methyl Imidazole Acetic acid | 0.38 | 1.28 | Down |
| 4 | C00262 | Hypoxanthine | 0.41 | 1.09 | Up/No change |
| 5 | C00750 | Spermine | Not detected |  |  |
| 6 | C00253 | Nicotinic acid | 0.38 | 1.28 | Down |
| 7 | C03824 | 2-Aminomuconic acid semialdehyde | 0.94 | 1.28 | Down |
| 8 | C10164 | Picolinic acid | 0.94 | 1.28 | Down |
| Downregulated<br>$p \leq 0.05$; $FC \geq 1.5$ | | | | | |
| 9 | C11695 | Anandamide | Not detected |  |  |
| 10 | C01753 | Beta-Sitosterol | 0.46 | 1.34 | Up |
| 11 | C04530 | 4,4-dimethyl-5alpha-cholest-7-en-3beta-ol | 0.46 | 1.34 | Up |
| 12 | C15915 | 4,4-Dimethyl-5a-cholesta-8-en-3b-ol | Not detected |  |  |
| 13 | C02528 | Chenodeoxycholic acid | Not detected |  |  |
| 14 | C04483 | Deoxycholic acid | Not detected |  |  |

Among the 14 metabolites detected in C vs LPS+ cells; only glutathione showed approaching significance with p value of 0.08, while for the other 7 metabolites, the change was non-significant but showed opposite trend whereas remaining 6 metabolites were not detected in LPS+ vs LPS+G+ comparison group (Table 2). Box plots for glutathione in both the comparison groups (C vs LPS+ &LPS vs LPS+G+) are shown in supplementary figure 2A and B.

### 3.5 Physiological relevance of differential metabolites in fatty liver diseases

LPS induced endotoxic shock in Caco-2 cells, resulting in significant upregulation of glutathione, hypoxanthine, spermine, nicotinic acid, 2-aminomuconic acid semialdehyde and picolinic acid; while anandamide, Beta sitosterol, Chenodeoxycholic acid and Deoxycholic acid and 4, 4-dimethyl-5alpha-cholest-7-en-3beta-ol or 4, 4-Dimethyl-5a-cholesta-8-en-3b-ol were downregulated in HepG2 cells as shown in Table 1.

#### 3.5.1 Metabolites with differential trends in LPS+ hepatocytes with or without G+

**Glutathione** (GSH), a tripeptide (G-Glutamate, S-S-Cysteine, H-Glycine) is a thiol reducing agent available at high concentration in liver, is observed to be upregulated due to endotoxic shock in hepatic cells ^31^. Moreover, serum levels of GSH were reported to be increased in patients diagnosed with non-alcoholic steatohepatitis (NASH) ^32^. In the present study, HepG2 cells in co-culture showed a significant two-fold (Table 1; p<0.05, FC=2.19) increase in glutathione levels compared to control HepG2 cells. LPS treatment to Caco-2 in co-culture, compromised the barrier function mimicking leaky gut and passage of LPS or LPS induced inflammatory metabolites to liver cells (HepG2) in the lower chamber of transwell plate, thus, mimicking condition of NASH in model of gut-liver axis. In the case of glucan pre-treatment followed by LPS, glutathione was down regulated by 1.44-fold (p=0.08). This may indicate protection by glucan against LPS-mediated liver injury.

**Uridine,** a pyrimidine nucleoside, was observed to be significantly increased by 1.66-fold in LPS treatment due to endotoxin-induced liver injury via disrupted gut-liver axis, thus impacting nucleotide metabolism ^33^. Uridine was not detected in glucan-pre-treated liver cells, showing no impact on pyrimidine metabolism in the presence of exopolysaccharide glucan.

**Methyl imidazole acetic acid (MIMA),** a histamine metabolite, was elevated as a consequence of LPS exposure, triggering inflammatory stimulus in HepG2 cells, but was down-regulated (FC= 1.28) in glucan pre-treated liver cells although not significant (p=0.38). LPS induced liver injury triggers histamine activity and metabolism that breaks down to release MIMA ^33–35^. Glucan pre-treatment reduced the levels of MIMA indicating role in preventing inflammation.

**Hypoxanthine**, a purine catabolite and substrate for the xanthine oxidase enzyme, was significantly upregulated (FC=2.31) in LPS-treated HepG2 cells. Studies in apolipoprotein E knock out mice (Apoe KO) and HepG2 cells have reported increased cholesterol accumulation via ROS generation due to hypoxanthine treatment ^36, 37^. Hence, the increased hypoxanthine levels could have contributed to hepatic steatosis in this gut-liver axis model. G+ co-treatment has brought down levels to one-fifth.

**Spermine**, a polyamine, is a ubiquitous small organic molecule found along with other polyamines, Putrescine and spermidine were observed to be upregulated in steatosis as a consequence of increased inflammation in the gut due to LPS release by dysbiotic gut flora ^38, 39^. Interestingly increased concentration of spermine is synthesized and secreted by gut microbes inhabitant of gut lumen, also in colonic epithelial cells as a consequence of endotoxin exposure ^40, 41^, for protecting the leaky gut lining and subsequent endotoxin spillage into systemic circulation, thus, suppressing inflammation. The studies on gut polyamine metabolism have shown their transport from intestinal epithelium via blood to various organs like liver, heart, and brain for beneficial effects ^42^. In the present study, we observed a 2.37-fold increase in spermine in LPS-treated HepG2 cells, which were co-cultured with Caco-2 cells, reflecting rapid renewal and proliferation of the colonic epithelial cells due to endotoxin exposure that needed increased uptake and metabolism of the polyamines ^40^. Spermine was not detected in the glucan co-treated HepG2 cells. This could be due to glucan neutralizing the inflammatory trigger by LPS that could have blocked enhanced spermine synthesis in caco-2 and subsequent transport to HepG2 cells, preventing liver cell injury. As per the literature microbial exopolysaccharides indirectly influence the spermine levels ^43^ ^42^; However, further experiments are to be carried out to confirm the involvement of glucan in polyamine transport to hepatic cells.

**Nicotinic acid (NA)**, niacin or vitamin B3, is reported to protect LPS-induced Caco-2 inflammation by lowering inflammatory cytokine ^44^, with pharmacological doses reported to have improved dyslipidemia by reducing triglycerides and LDL cholesterol while raising HDL cholesterol ^45^. But NA supplementation as lipid lowering drug is discontinued due to its side effects, including hyperglycemia and hyperuricemia, also raising liver aminotransferases ^46^. Additionally, studies showed, higher doses can be detrimental to liver and inhibitors of NA to be beneficial ^47^. However, in a recent cohort study from National Health and Nutrition Examination Survey (NHANES) on the US population published patients with NAFLD may benefit from a higher intake of dietary niacin. ^48^. Therefore, there were contradictory reports regarding the effects of niacin on liver health. In our study, we observed a significant 4.21-fold increase in NA in LPS-treated pathological state compared to control liver cells co-cultured with colonocytes, and it was downregulated to 1.28-fold in glucan pretreatment. In contrast, in our analysis, we had observed a seven-fold increase in niacinamide, the amide of nicotinic acid, in glucan-pretreated LPS+ liver cells. Thus, indicating glucan as a modulator of niacin levels could play a pivotal role in LPS-induced liver injury.

**2-Aminomuconic acid semialdehyde and Picolinic acid** are metabolites of the oxidative metabolism of tryptophan via the kynurenine pathway in mammalian cells. 2-Aminomuconic acid semialdehyde is unstable and spontaneously converted to picolinic acid ^49^. Kynurenine pathway is the main catabolic pathway, and the enzyme tryptophan 2, 3-dioxygenase (TDO) is highly expressed in liver, and is one of the rate-limiting steps that converts tryptophan to N-formylkynurenine ^50^. Increased tryptophan metabolism via the kynurenine pathway is associated with an increase in inflammation and lipid accumulation in the liver, resulting in NAFLD ^49^. In our analysis, there was a significant 4.21-fold increased detection of metabolites 2-aminomuconic acid semialdehyde and picolinic acid in LPS-treated HepG2 cells, indicating proinflammatory and lipotoxic liver injury. Both of these metabolites were down-regulated by 1.28-fold in glucan treatment. Thus, glucan affecting tryptophan metabolism via the kynurenine pathway could protect against hepatic steatosis, preventing progression to NAFLD or NASH. Thus, these metabolites could be interesting as prognostic markers in glucan-mediated therapy on liver injury in future studies.

**Beta-Sitosterol (BS)**, a phytosterol found in plenty in plants, has been reported to prevent NAFLD by reducing oxidative stress, inflammatory responses, as well as regulating fatty acid oxidation factors such as proliferator-activated receptor (PPAR-α), sterol regulatory element binding protein (SREBP-1c), and carnitine palmitoyltransferase-1(CPT-1) ^51, 52^. In the present study, BS was significantly down by 1.73-fold in LPS alone-treated liver cells, while it was up by 1.34-fold in glucan co-treated liver cells. Thus, it could be one of the positive biomarkers to identify the glucan-mediated effect.

Similar to BS, 4, 4-dimethyl-5alpha-cholest-7-en-3beta-ol, an intermediate in cholesterol metabolism, was 1.73-fold down-regulated in LPS-treated HepG2, which was up by 1.34-fold in glucan-pretreated liver cells.

#### 3.5.2 Other significantly changed metabolites detected only in LPS+ compared to no treatment cells

Primary bile acids like Chenodeoxycholic acid (CDCA) and Deoxycholic acid (DCA) were significantly downregulated by 1.62-fold in LPS+ liver cells compared to control. Primary bile acids play a key role in digestion, cholesterol metabolism and absorption ^53^. Lower levels of bile acids result in cholesterol accumulation in the liver, thus resulting in the condition of hepatic steatosis. Hence, the LPS-treated co-culture model of gut-liver axis resulted in fat deposition in HepG2 cells via cholesterol accumulation. However, these metabolites were not detected in LPS+G+ cells.

### 3.6 Metabolomics pathway enrichment and impact analysis

KEGG Mapper for pathway mapping was used to obtain enrichment of annotated metabolites having KEGG identifiers in metabolic pathways. The mapping suggested metabolite hits in 56 metabolic pathways of Homo sapiens (human) database (selected library code ‘hsa’), in LPS+ vs LPS+G+ comparison group. Among the hit pathways, the maximum metabolite hits were of steroid hormone biosynthesis pathways (n=35 hits) and primary bile acid biosynthesis pathways (n=17 hits) (Figure 5A). Pathway impact (PI) assessed the impact of different pathways in LPS (LPS+) challenged hepatic cells in response to glucan treatment (LPS+G+) as shown in Figure 5B. The centrality and pathway enrichment results were combined to measure the high-impact pathways. The most significant (0.05) and highly enriched pathway having high impact in glucan treatment to LPS challenged liver cells included those involved in biosynthesis pathways of steroid hormone (PI=0.46), primary bile acid (PI=0.27), Phenylalanine, tyrosine, and tryptophan (PI=0.5), and steroid (PI=0.3). Whereas Galactose (PI=0.78), linoleic acid (PI=1), starch and sucrose (PI=0.55), sphingolipid (PI=0.44), arginine and proline (PI=0.28), tyrosine (PI=0.15), amino and nucleotide sugar (PI=0.14), as listed in supplementary table 2. The top 25 enriched metabolic pathways were shown (Figure 5C) following enrichment analysis by Metabolite Set Enrichment Analysis (MSEA) in MetaboAnalyst 5.0. The analysis results showed steroid hormone biosynthesis as the topmost and primary bile acid biosynthesis to be in the top 3 position. Enriched pathway of steroid hormone synthesis indicated an increase in cholesterol biosynthesis and accumulation. As cholesterol deposition was reported in steatosis conditions as a result of increased gut dysbiosis and its impact on steroid hormone biosynthesis ^4^.

**Figure 5.**
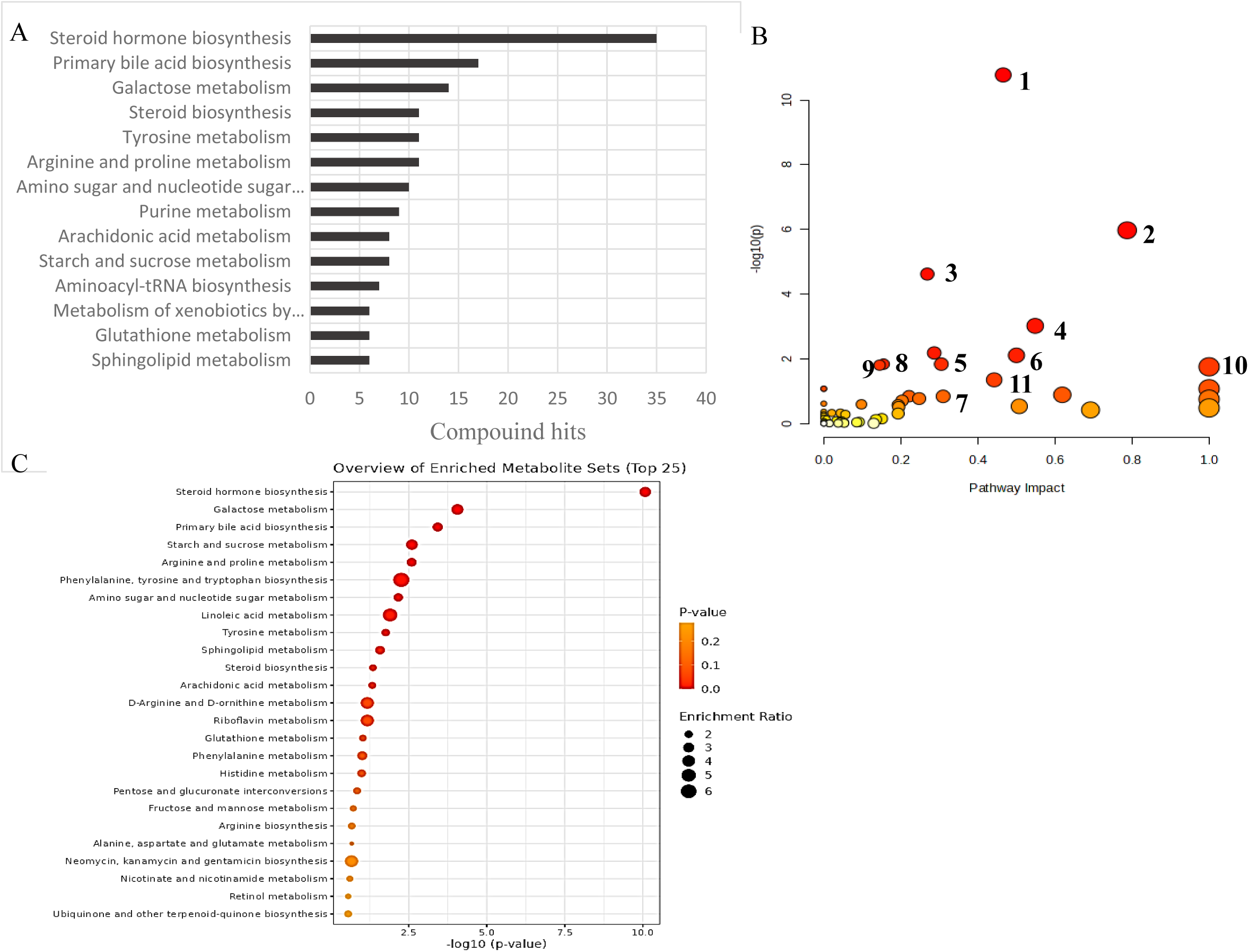
Pathway enrichment analysis using KEGG Mapper mapped metabolite feature hits with different biosynthesis and metabolic pathways in LPS+ vs LPS+G+. (Compound hits >5 no.s are shown) (A) Impact of significantly different pathways (p≤0.05) are shown by pathway impact analysis in caco-2 co-cultured liver cells exposed to LPS with or without glucan treatment; (B) Color of circle from red to yellow represent high to low significance levels while size of the circle indicates the impact of the pathway, identified pathways were (1) Steroid hormone biosynthesis (2) Galactose metabolism (3) Primary bile acid biosynthesis (4) Starch and sucrose metabolism (5) Phenylalanine, tyrosine and tryptophan biosynthesis (6) Arginine and proline metabolism (7) Steroid biosynthesis (8) Tyrosine metabolism (9) Amino-sugar and nucleotide sugar metabolism (10) Linoleic acid metabolism (11) Sphingolipid metabolism; (C) Metabolite Set Enrichment Analysis (MSEA) showed top 25 enriched pathways based on functionally related metabolite sets.

Biosynthesis of primary bile acids in the liver is responsible for the disposal of cholesterol from systemic circulation. ^4, 54^. The significantly increased primary bile acid levels may indicate the effect of LPS-induced liver steatosis condition affecting liver toxicity and inflammation. Further analysis of individual metabolites in the bile acid biosynthesis pathway by experimental analysis is required to validate these findings.

## 4. Conclusions

The current study demonstrates testing of glucan in hepatic steatosis in a cell culture model of the gut-liver axis. To mimic gut dysbiosis and establish liver steatosis condition in liver cells, HepG2 cells were co-cultured with Caco-2 cells in transwell plates and treated with pathological doses of LPS at 10µg/ml for 24 hours. To study the effect of glucan, pretreatment of 10 mg (weight equivalent of unfermented glucan) fecal fermented glucan for 24 hours was carried out to compare with LPS alone-treated cells. We observed an increase in lipid deposition determined by Oil Red O staining and altered gene expression associated with lipid metabolism, viz. HMGCR (increased) and ACCB (reduced) as quantified by RT-PCR. After confirmation of the steatosis model, metabolomics of the treated cells was carried out to analyze the differential metabolomic profiles of the liver cells as a result of treatment.

Untargeted metabolomics of the intracellular metabolites of HepG2 cells was carried out using high-resolution mass spectrometry (LC-MS/MS). Metabolomic profiles of LPS+ with no treatment control and LPS+ with glucan co-treated LPS+ hepatic cells were compared employing advanced statistics. We observed significantly upregulated metabolites, viz. glutathione, uridine, methyl imidazole acetic acid (MIMA), hypoxanthine, spermine, nicotinic acid (NA), 2-amino muconic acid semialdehyde, and picolinic acid which were associated with liver steatosis via mechanism of lipid accumulation, oxidative stress and inflammation in a thorough literature analysis. Except for MIMA and NA, all other metabolites were down-regulated in glucan co-treated liver cells. Other metabolites such as beta sitosterol and 4, 4-dimethyl-5alpha-cholest-7-en-3beta-ol were down-regulated in LPS treated liver cells, but in glucan pre-treatment, these were found to be up-regulated. Although threshold of fold change was reversed, it was not significant, probably due to the small sample size. Additionally, pathway analysis and pathway impact showed a significant number of hits primarily related to steroid hormone biosynthesis and primary bile acid synthesis pathways in both LPS and glucan-treated conditions. This indicated that cholesterol metabolism is the primary target of glucan affected in the liver due to steatosis. Hence, our cell culture model of gut-liver axis potentially indicated a modulating effect of glucan on biochemical pathways of metabolites associated with liver injury, especially cholesterol metabolism due to gut dysbiosis, to reverse hepatic steatosis.

## Limitations of the study

The sample size for cell culture-based metabolome evaluation was carried out on three biological samples with three technical replicates.

## Author contribution

Study design, conducted experiments, data analysis, writing original draft editing, review-URP; Conducted experiments, data analysis, editing, review-NG; contributed to material-ST and IAS; Supervision, data acquisition, contributed to reagents/materials/analysis tools, data analysis, editing, review –DKP; Original idea, supervision, study conception, data acquisition, contributed to reagents/materials/analysis tools, data analysis, editing, review-PHS; Conflict of interest: The authors declare no competing interest

## Data availability

On request

## Acknowledgement

URP acknowledges ICMR, India, for RA fellowship (file no. 2021-9505/CMB-BMS). URP acknowledges Palanisamy Bruntha Devi for technical advise on fecal fermentation of glucan.

---

**Supplementary Figure 1.**
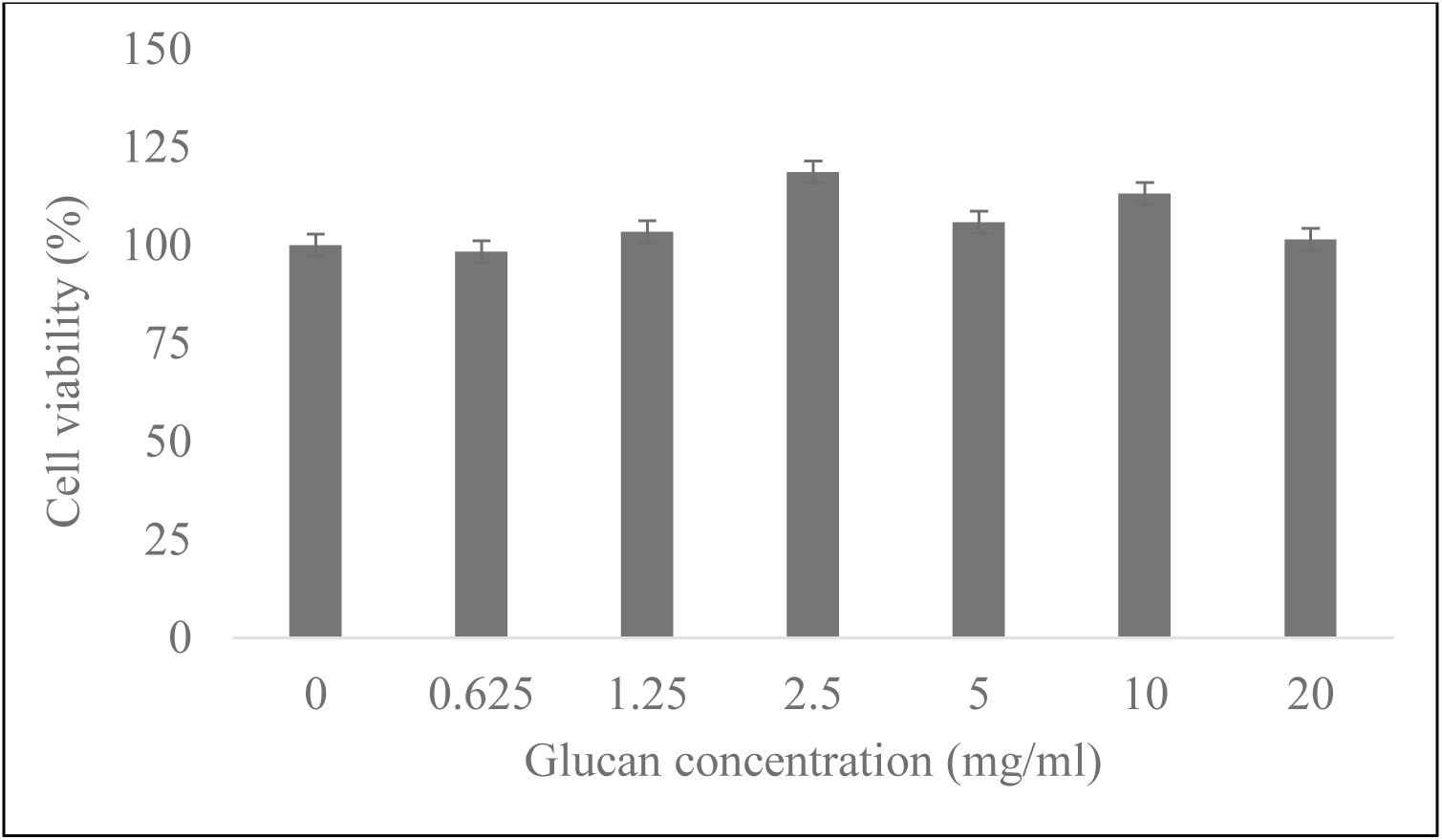
Cell viability of HepG2 in co-culture with Caco-2 after 24 hours’ treatment with fermented glucan at concentrations of 0, 0.625, 1.25, 2.5, 5, 10, and 20 mg/ml weight equivalent of non-fecal fermented glucan by MTT assay. All % cell viability values were expressed as mean ±SEM of 3 replicates.

**Supplementary figure 2.**
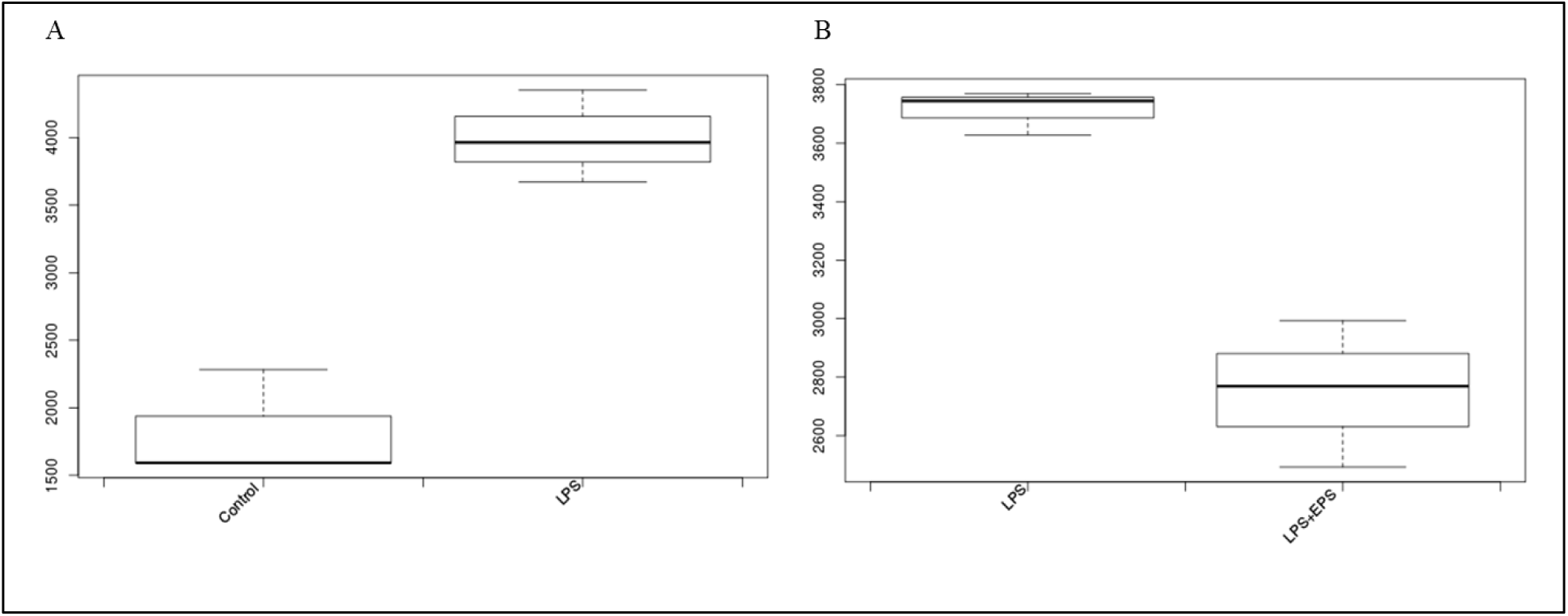
Boxplot of Glutathione in two comparison groups. (A) There was significant (p=0.002) 2.2-fold increase in LPS+ treated cells compared to control; (B) It was 1.44-fold down regulated (p=0.08) in glucan/ fermented EPS co-treated (LPS+EPS/G+) cells compared to LPS+ alone treatment.

**Supplementary Table 1.**
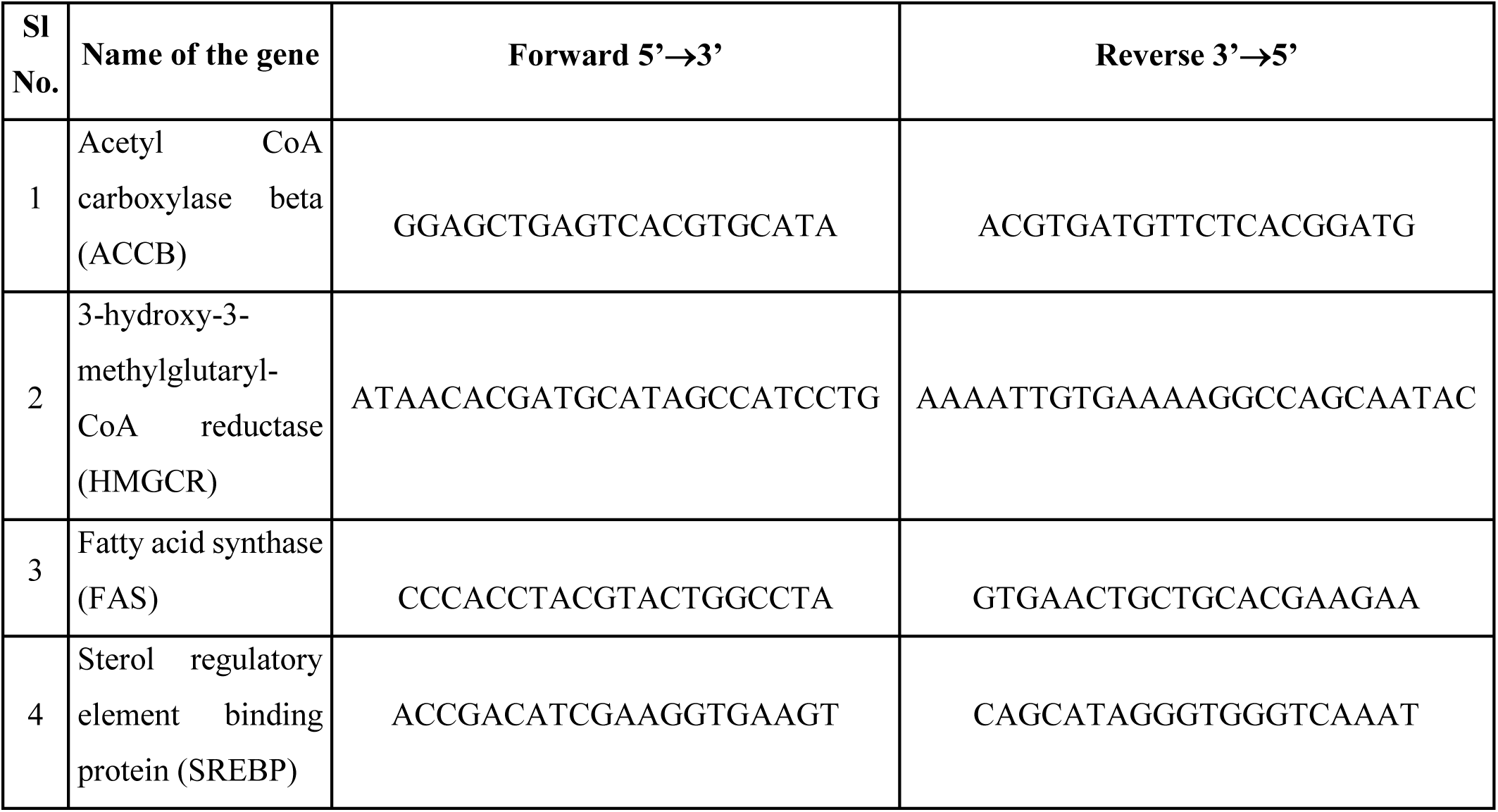
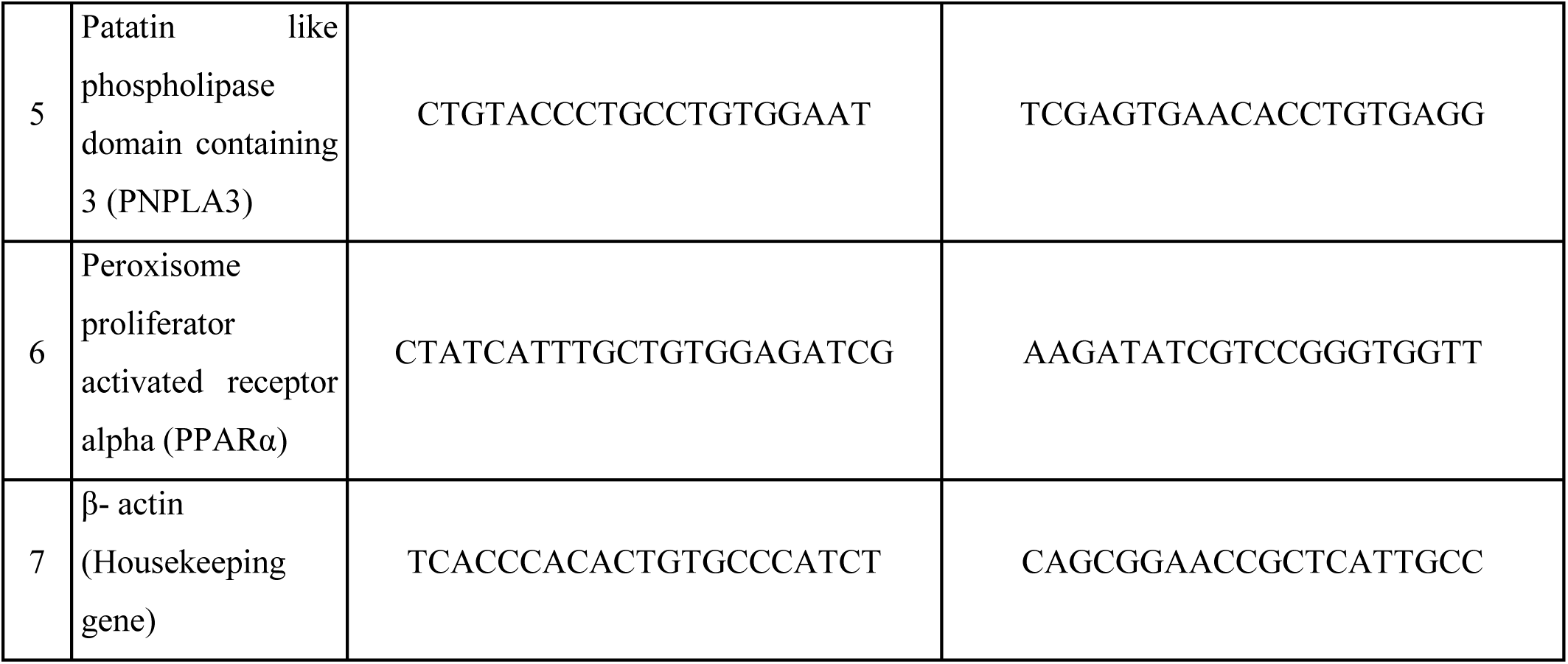
Sequence of primers used in quantitative real-time PCR experiments.

**Supplementary Table 2:** Pathways having significantly high impact in LPS+ compared with LPS+G+ treated cells.

| Sl no. | Pathway name | p value | Pathway impact (PI) |
| --- | --- | --- | --- |
| 1 | Linoleic acid metabolism | 0.017602 | 1.000 |
| 2 | Galactose metabolism | 1.06E-06 | 0.787 |
| 3 | Starch and sucrose metabolism | 0.00095718 | 0.549 |
| 4 | Phenylalanine, tyrosine and tryptophan biosynthesis | 0.0077832 | 0.500 |
| 5 | Steroid hormone biosynthesis | 1.67E-11 | 0.466 |
| 6 | Sphingolipid metabolism | 0.0446 | 0.442 |
| 7 | Steroid biosynthesis | 0.014565 | 0.305 |
| 8 | Arginine and proline metabolism | 0.0065629 | 0.287 |
| 9 | Primary bile acid biosynthesis | 2.39E-05 | 0.269 |
| 10 | Tyrosine metabolism | 0.014565 | 0.155 |
| 11 | Amino sugar and nucleotide sugar metabolism | 0.015661 | 0.145 |

## Notes

### Competing Interest Statement

The authors have declared no competing interest.

